# Robustness of early warning signals for catastrophic and non-catastrophic transitions

**DOI:** 10.1101/218297

**Authors:** Partha Sharathi Dutta, Yogita Sharma, Karen C. Abbott

## Abstract

Early warning signals (EWS) are statistical indicators that a rapid regime shift may be forthcoming. Their development has given ecologists hope of predicting rapid regime shifts before they occur. Accurate predictions, however, rely on the signals being appropriate to the system in question. Most of the EWS commonly applied in ecology have been studied in the context of one specific type of regime shift (the type brought on by a saddle-node bifurcation, at which one stable equilibrium point collides with an unstable equilibrium and disappears) under one particular perturbation scheme (temporally uncorrelated noise that perturbs the net population growth rate in a density independent way). Whether and when these EWS can be applied to other ecological situations remains relatively unknown, and certainly underappreciated. We study a range of models with different types of dynamical transitions (including rapid regime shifts) and several perturbation schemes (density-dependent uncorrelated or temporally-correlated noise) and test the ability of EWS to warn of an approaching transition. We also test the sensitivity of our results to the amount of available pre-transition data and various decisions that must be made in the analysis (i.e. the rolling window size and smoothing bandwidth used to compute the EWS). We find that EWS generally work well to signal an impending saddle-node bifurcation, regardless of the autocorrelation or intensity of the noise. However, EWS do not reliably appear as expected for other types of transition. EWS were often very sensitive to the length of the pre-transition time series analyzed, and usually less sensitive to other decisions. We conclude that the EWS perform well for saddle-node bifurcation in a range of noise environments, but different methods should be used to predict other types of regime shifts. As a consequence, knowledge of the mechanism behind a possible regime shift is needed before EWS can be used to predict it.

## Introduction

Absent any large and sudden perturbations, we do not typically expect ecosystems to exhibit large and sudden changes in their state. Exceptions to this rule are the subject of major concern because they represent the alarming situation where small external changes create dramatic shifts in the composition, configuration, and possibly function of an ecosystem (Holling 1973, Scheffer 2009). Such sudden regime shifts can occur in two main ways (Shea et al 2004). First, a small perturbation to a parameter, such as a demographic rate or interaction term, may cause a system to cross a bifurcation point – a critical point beyond which the qualitative dynamics of the system change. Second, a small perturbation to the system state may push a system with multiple stable configurations into the basin of attraction of a different stable state. In either case, the small external changes that trigger the shift are effectively cryptic and recognizing when a particular system is at risk of suddenly shifting to a different regime is a formidable challenge.

A growing body of research on early warning signals (EWS) has recently been developed to meet this challenge. EWS are statistics associated with the detection of rapid regime shifts before they occur. EWS have been developed for a diverse range of dynamical scenarios, e.g. critical transitions in ecosystems (Scheffer et al 2009), onset of neuron spiking (Meisel et al 2015), rate-induced tipping in climate system (Ashwin et al 2012, Ritchie and Sieber 2016, 2017), and transitions in non-stationary models (Kwasniok 2015) and networks (Mheen et al 2013, Kuehn et al 2015). However, EWS in ecology have mostly been used to detect a specific type of regime shifts where the current state of a system is one of two stable states, and where incremental external changes will soon cause the system to cross a bifurcation (specifically a saddle node, also known as a fold, bifurcation) where the current state and an unstable equilibrium point merge and disappear. Loss of the current stable state will force the system to shift to the other stable state. A stable equilibrium is characterized by a positive rate of return following a local perturbation, and this rate approaches zero as the system nears a bifurcation at which the equilibrium vanishes or loses stability (Wissel 1984). This phenomenon in which return rates approach zero, known as critical slowing down (Strogatz 1994, Van Nes and Scheffer 2007), has certain generic effects on dynamics like increased variance and autocorrelation near bifurcations; we therefore expect to see these effects when the current state of a system is about to cease being stable (Scheffer et al 2009, Held and Kleinen 2004, Brock and Carpenter 2006, Carpenter and Brock 2006, Kleinen et al 2003, Guttal and Jayaprakash 2008, Seekell et al 2011). Although EWS have been successfully applied in some cases (Scheffer et al 2009), a thorough understanding of when and how they work reliably is still an open area of research (Boettiger and Hastings 2012b, Drake 2013, Boettiger and Hastings 2013, Boettiger et al 2013, Dakos et al 2015, Gsell et al 2016).

As eloquently mapped out in Boettiger et al (2013), rapid regime shifts may or may not involve bifurcations, and bifurcations may occur with or without critical slowing down (CSD). Because EWS are generic symptoms of CSD, rather than being indicators of regime shifts per se (Van Nes and Scheffer 2007, Kéfi et al 2012), they tend to work well when regime shifts and CSD co-occur. This is the situation with some catastrophic bifurcations (“catastrophic” meaning those with a large qualitative effect not readily reversed) like the saddle node. However, when (a) regime shifts occur without CSD, or (b) CSD occurs without an associated regime shift, EWS become more difficult to interpret. Situation (a) arises for catastrophic bifurcations that lack CSD (e.g. Hastings and Wysham 2010, Schreiber and Rudolf 2008) or when regime shifts are due to stochastic switching between coexisting stable states in the absence of any external change that would trigger CSD (e.g. Boettiger and Hastings 2012a, Sharma et al 2015). For these transitions, EWS are not expected (Hastings and Wysham 2010, Boettiger and Hastings 2013). Situation (b) occurs for non-catastrophic transitions like super-critical Hopf, transcritical, and pitchfork bifurcations, that have CSD but are characterized by quantitatively similar dynamics before and after bifurcation. For example, on one side of a super-critical Hopf bifurcation is a stable node and on the other side is a limit cycle, initially very small, that is centered around that node. Although there is a meaningful qualitative change of dynamics at the bifurcation (transition from a point equilibrium to a cycle), the actual change in population densities or ecosystem state at the bifurcation point is trivial. Larger cycles farther past the bifurcation point can certainly be ecologically important, but the key here is that at the bifurcation point itself, there is no meaningful regime shift. Nevertheless, because these bifurcations occur with CSD, EWS can appear (Kéfi et al 2012).

In addition to transition type – catastrophic or not, with or without a bifurcation or CSD – there are important open questions about how noise type affects the performance of EWS (Contamin and Ellison 2009). The “slowing down” of CSD refers to a system’s rate of recovery following a perturbation. EWS thus only appear because perturbations are present, and these are usually in the form of stochastic noise. The properties of this noise are likely to have an effect on statistics like population variance and autocorrelation. For instance, positively autocorrelated (red-shifted) noise can have a substantial impact on population dynamics (Ripa and Lundberg 1996, and others), and how this effect interacts with the changes in population variance and autocorrelation caused by critical slowing down is not fully understood. Rudnick and Davis (2003) found that noise color strongly influenced the performance of a regime shift indicator in purely stochastic time series, but neither Perretti and Munch (2012) nor Boerlijst et al (2013) saw an effect of noise color on the performance of EWS in stochastic population models. This led Perretti and Munch (2012) to hypothesize that noise color may have a stronger effect on EWS when population dynamics are more strongly influenced by noise, but to our knowledge this idea has never been tested. Higher noise intensity could cause population dynamics to be more strongly influenced by noise, but so too could CSD itself, where a slower recovery from perturbation means weaker intrinsic regulation. A clearer understanding of if and when noise color affects the performance of EWS is needed (Boettiger and Hastings 2012a), especially given the commonness of red-shifted noise in ecological systems (Halley 1996, Vasseur and Yodzis 2004). Aside from noise color, other properties, like whether perturbations cause variation in parameter values, or to total or per-capita population growth rates, may also affect EWS performance (Dakos et al 2012b).

Finally, when computing EWS from data, researchers must make decisions about how much data before a transition to use and over what rolling window with what smoothing bandwidth. Examples have shown EWS to be sensitive to time series length (Sharma et al 2015), sampling interval (Perretti and Munch 2012), and, in some instances, bandwidth and window size (Dakos et al 2012a). However, the extent to which these individual results hold across different transition types and with different types of noise has not yet been explored.

In this paper, we follow Kéfi et al (2012) and compute two major early warning signals, variance and lag-1 autocorrelation, for a suite of models with different transition types. To complement Kéfi et al (2012), who used additive white (uncorrelated) noise in their models, we consider multiplicative white and red-shifted (positively autocorrelated) noise. Our models span all 5 dynamical categories delineated by Boettiger et al (2013) (redrawn in figure 1a): (I) the “charted territory” of the saddle node bifurcation, with CSD and a regime shift; and the “uncharted territories” of (II) regime shifts through bifurcations that lack CSD; (III) non-catastrophic bifurcations with CSD; (IV) CSD without a bifurcation; and (V) regime shifts due to stochastic switching, absent a bifurcation and CSD. We also computed EWS for a model undergoing a smooth and gradual state change, with no regime shift, bifurcation, or CSD, as a null case. For each model, we considered a range of time series lengths, rolling window sizes, and bandwidths in our calculations, to determine the effect of these choices on EWS performance. As expected, we find that EWS are more sensitive to time series length in comparison with window size or bandwidth within the ranges we tested. Although EWS work as expected (provided a sufficiently long time series) in the case of the saddle node bifurcation, we find both surprising positive and surprising negative results for other transition types. Because Kéfi et al (2012) did not see similar surprises when examining many of the same transitions using additive white noise, we conclude that EWS performance is highly sensitive to noise type. Together, our results reemphasize the need for the mechanisms underlying a possible regime shift to be understood first, before EWS can be applied and properly interpreted (Boettiger et al 2013).

**Figure 1:**
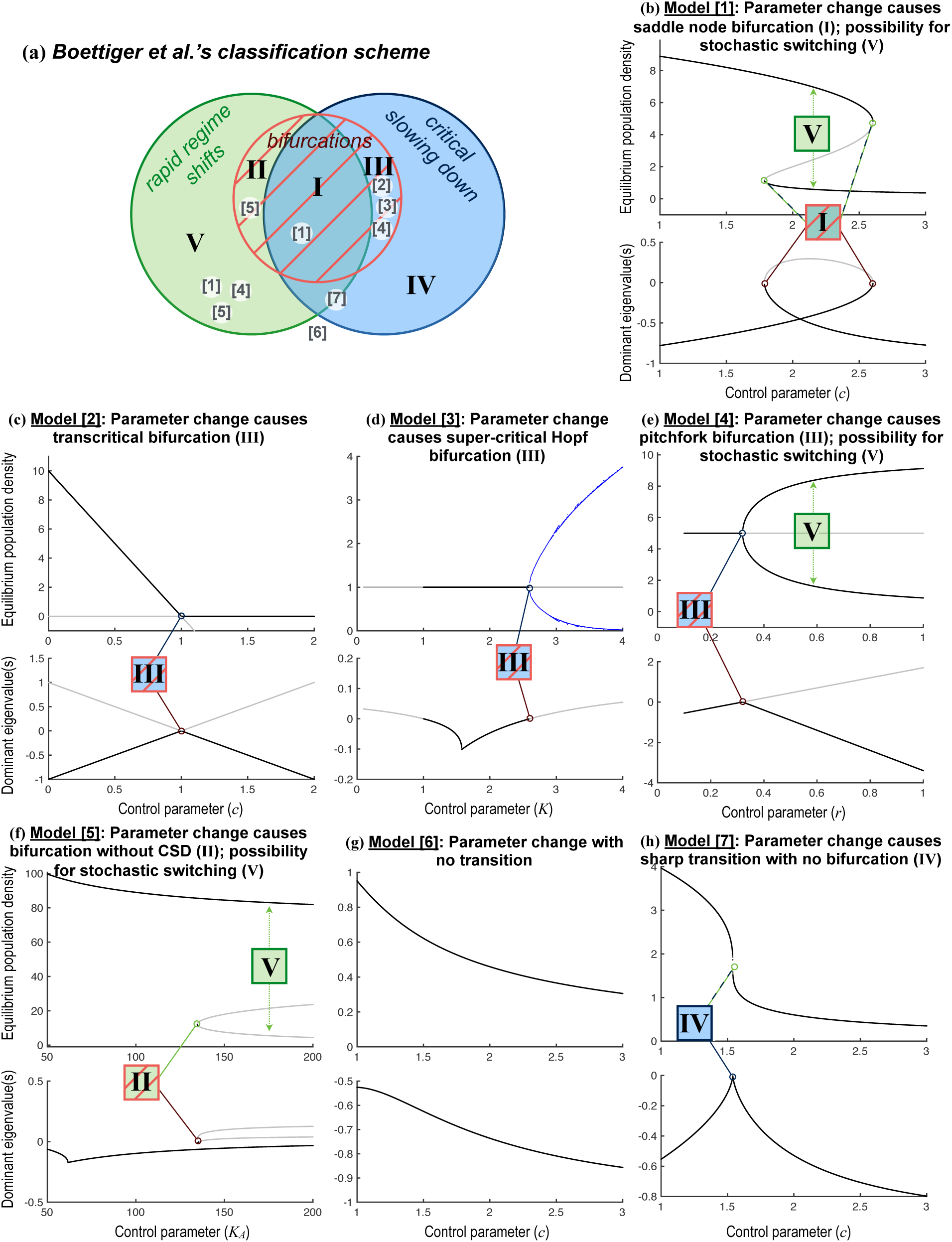
(a) Venn diagram, redrawn and slightly modified from Boettiger et al (2013), showing possible combinations of critical slowing down, rapid regime shifts, and bifurcations. Roman numerals label the 5 distinct combinations, as laid out in Boettiger et al (2013); we modified region IV slightly to include any case of critical slowing down that occurs in the absence of a bifurcation (regardless of whether or not a rapid regime shift also occurs; this allows us to place the sharp transition in (h) cleanly into region IV without debating whether the transition truly qualifies as a rapid regime shift). Numbers in square brackets correspond to the model numbers used in table 1 and throughout, showing where each model fits within this classification scheme. (b-h) Bifurcation diagrams and dominant eigenvalues (real part) for each model and parameter set shown in table 1. Stable equilibria (upper graph in each pair) and their dominant eigenvalues (lower graph) are plotted in black; unstable equilibria and their dominant eigenvalues are plotted in gray. Transitions are labeled using the roman numerals from (a). Beyond the super-critical Hopf bifurcation in model [**3**] (panel (d)), we show the amplitude of limit cycles in blue. **Bifurcations** occur in panels (b)-(f) where eigenvalues intersect 0. **Critical slowing down** occurs as the bifurcation is approached in (b)-(e), and when the eigenvalue peaks sharply just below 0 in (h). **Rapid regime shifts** are possible due to changes in the control parameter in (b), (f), and (h), where the state before and after transition is quite different. Rapid regime shifts can also occur due to stochastic switching in (b) and (e) where 2 stable point equilibria coexist for a given value of the control parameter, and in (f) where the upper point equilibrium coexists with a limit cycle around the lowest unstable equilibrium (Schreiber and Rudolf 2008).

## Methods

### Simulations

We begin by reanalyzing the models studied by Kéfi et al (2012) for a different noise type (multiplicative noise, as described below). These models include a well-known example of a saddle node bifurcation (Noy-Meir 1975, May 1977, Ludwig et al 1978) (table 1 model [**1**], figure 1b) and a well-known model with a super-critical Hopf bifurcation (Rosenzweig 1971) (table 1 model [**3**], figure 1d), as well as modified versions of model [**1**] that instead have a transcritical (table 1 [**2**], figure 1c) or pitchfork (table 1 [**4**], figure 1e) bifurcation. Also following Kéfi et al (2012), we consider model [**1**] with two alternative parameter settings: one in which the model lacks a transition altogether (the null case: table 1 [**6**], figure 1g) and one that produces an eigenvalue peak (and thus relative slowing of recovery from perturbations) but no bifurcation (table 1 [**7**], figure 1h). To this list, we add a model that exhibits a bifurcation and regime shift without CSD (Schreiber and Rudolf 2008) (table 1 [**5**], figure 1f).

**Table 1.**
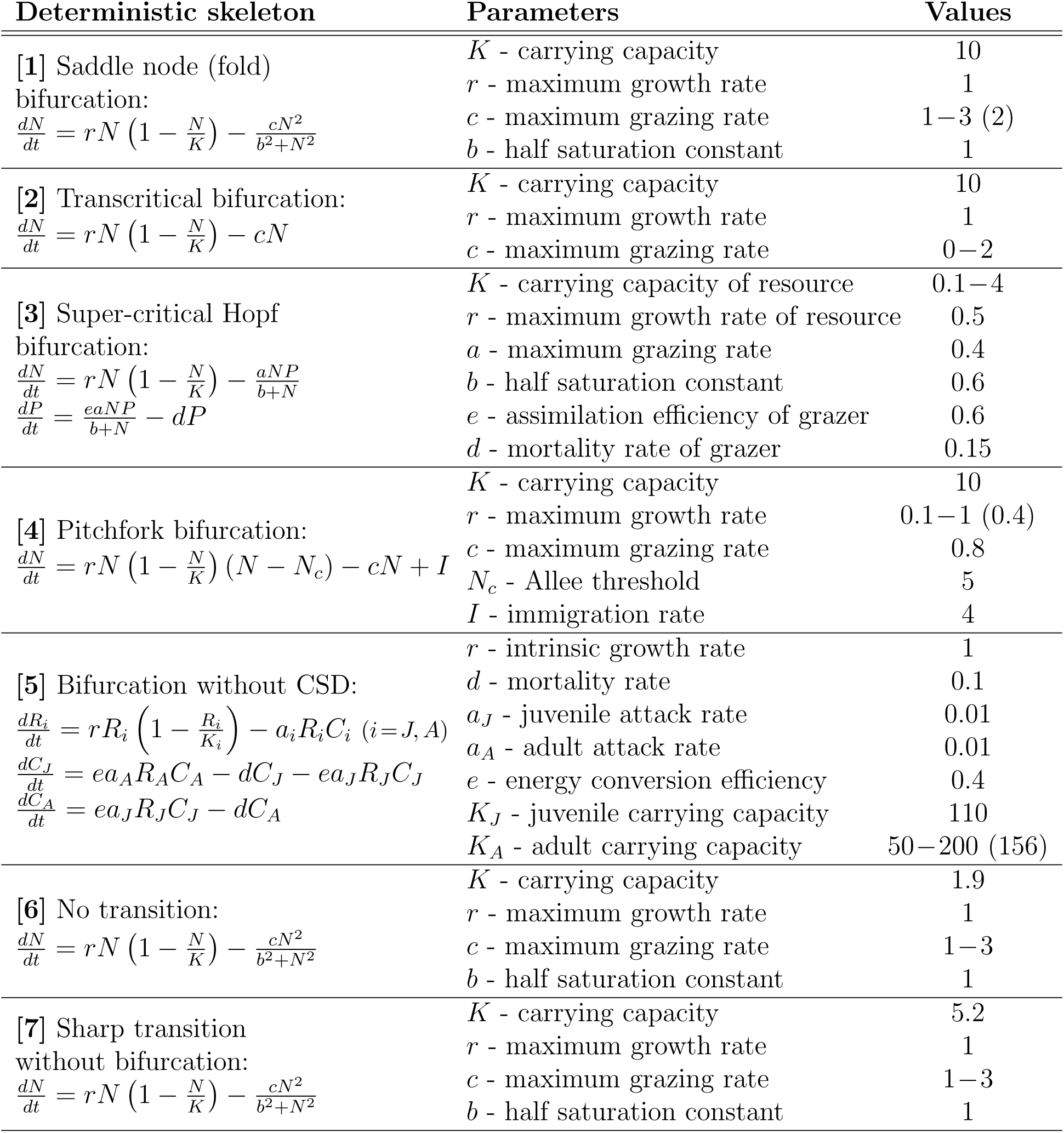
Deterministic base models, parameters descriptions, and values used. For each model, the control parameter is shown with a range of values. To move the system across the bifurcation or other transition point, we gradually varied the control parameter across this range. To explore stochastic switching, we fixed all parameters including the control parameter; the fixed value used for the control parameter is shown in parentheses.

To each of these models, we incorporated multiplicative, red-shifted Gaussian noise representing autocorrelated environmental stochasticity. The stochastic models each have the general form,

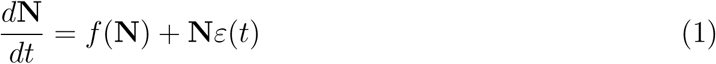

where N is population density (for one-dimensional models) or a vector of population densities, *f*(N) is the deterministic skeleton of the model as shown in table 1, and *ε*(t)is a random variable representing the colored noise. We refer to this formulation “multiplicative noise” because *ε*(t) is multiplied by a function (here, simply N) of population density. Kéfi et al (2012) used a common alternative, so-called “additive noise,” in which the random variable is added to the deterministic skeleton: 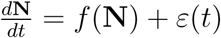. Additive noise represents random perturbations whose impact is independent of population size; random density-independent immigration and emigration are examples. Multiplicative noise represents perturbations with a per-capita effect, such as random fluctuations in survivorship or fecundity.

We write ε(t) in Eq. (1) as an Ornstein-Uhlenbeck process with derivative,

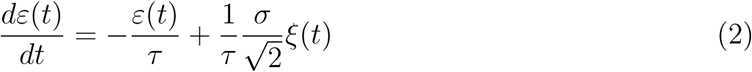

where *ξ* is the Gaussian white noise with zero mean and unit variance, *σ* is the noise intensity, and *τ* is the correlation time of the Ornstein-Uhlenbeck process. The autocorrelation function for *ξ* is,

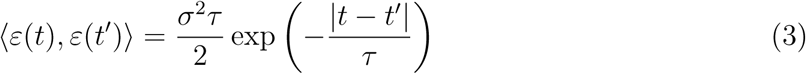

Although we focus on red-shifted noise (*τ* > 0) in this article, we briefly also consider the case of white noise, which occurs when *τ* → 0.

All of these models (except for the null case, model [**6**]) exhibit a transition of some sort as one “control parameter” is changed within the range shown in table 1. For models [**1**]-[**5**], these transitions are different kinds of bifurcations, and for model [**7**] the transition is a sharp but continuous change in the state variable (figure 1f). As the control parameter is changed, regime shifts will occur for catastrophic bifurcations ([**1**] and [**5**]) and in model [**7**], where the transitions involve a meaningful quantitative change in state, but not for the non-catastrophic bifurcations ([**2**]-[**4**], where the transition is only qualitative) nor for [**6**]. CSD occurs when eigenvalues approach 0, which happens when a local bifurcation is approached ([**1**]-[**4**]) or when the eigenvalue peaks sharply just below 0 ([**7**]; figure 1h).

For a fixed value of the control parameter, rapid regime shifts can also occur due to stochastic switching when multiple stable equilibria are present. This happens in models [**1**] and [**4**], where 2 stable point equilibria coexist for some values of the control parameter, and in [**5**] where the upper point equilibrium coexists with a lower limit cycle (Schreiber and Rudolf 2008) (figure 1b,e,f).

For each model, we performed stochastic simulations in MATLAB (R2011a) using the Euler-Maruyama method (Higham 2001) with standard integration step size of 0.001. We chose to study these models via simulation, both to mimic the way time series data are analyzed and to allow consistent treatment of the 1-dimensional models (for which one could study the dynamics by deriving and solving the master equation (H¨anggi and Jung 1995)) and models with higher dimension. Where our suite of models overlaps with Kéfi et al (2012), we used the same parameter values as they did to facilitate comparison. To examine transitions due to changes in the control parameter, we fixed all other parameters at the values shown in table 1 and simulated the model while gradually changing the control parameter across the range shown over 1000 time units. To examine stochastic switching, we fixed the control parameter at the value given in parentheses in table 1.

We simulated all models across a range of red-shifted noise environments by using various combinations of *σ* and *τ*, as given in the “Ranges” columns of table 2. We then followed up with one specific *σ − τ* combination for each model (“Fixed values” in table 2) for an in-depth comparison of the different window sizes, time series lengths, and bandwidths.

**Table 2.** Values of *τ* and *σ* used. ΔParam refers to simulations where the control parameter was changed to drive transitions and Stoch refers to simulations of stochastic switching. We ran each model under approximately 400 combinations of *σ* and *τ* chosen to span the range of behaviors observed in the parameter ranges given; these results are reported in aggregate as histograms. Specific examples we studied use the fixed values given in the last 2 columns.

| Model | Cause of transition | *Ranges* |  | *Fixed values* |  |
| --- | --- | --- | --- | --- | --- |
| | | $\sigma$ | $\tau$ | $\sigma$ | $\tau$ |
| [1] | ΔParam | $[3, 5] \times 10^{-5}$ | $[0.01, 0.03]$ | $3 \times 10^{-5}$ | 0.01 |
| [1] | Stoch | $[5, 9] \times 10^{-4}$ | $[0.01, 0.03]$ | $5 \times 10^{-5}$ | 0.01 |
| [2] | ΔParam | $(0, 8] \times 10^{-4}$ | $[0.1, 1.9]$ | $4 \times 10^{-4}$ | 1 |
| [3] | ΔParam | $[1, 9] \times 10^{-4}$ | $[0.1, 0.3]$ | $5 \times 10^{-4}$ | 0.1 |
| [4] | ΔParam | $[4, 8] \times 10^{-5}$ | $[0.1, 0.3]$ | $6 \times 10^{-5}$ | 0.1 |
| [4] | Stoch | $[1.4, 1.6] \times 10^{-4}$ | $[0.1, 0.3]$ | $1.5 \times 10^{-4}$ | 0.1 |
| [5] | ΔParam | $[3, 5] \times 10^{-3}$ | $[0.1, 0.3]$ | $4.2 \times 10^{-3}$ | 0.1 |
| [5] | Stoch | $[6, 8] \times 10^{-3}$ | $[0.01, 0.03]$ | $7 \times 10^{-3}$ | 0.01 |
| [6] | ΔParam | $[2, 4] \times 10^{-5}$ | $[0.01, 0.03]$ | $3 \times 10^{-5}$ | 0.01 |
| [7] | ΔParam | $[0.1, 1] \times 10^{-3}$ | $[0.1, 0.3]$ | $1 \times 10^{-4}$ | 0.1 |

Finally, because our results under multiplicative red-shifted noise differ notably from those of Kéfi et al (2012) under additive white noise, we repeated our analysis using multiplicative white noise (i.e. equation (1) with temporally uncorrelated *ε*(t), *τ* → 0). This allows us to assess the effect of red-shifted noise independently from our use of multiplicative perturbations.

### Measuring EWS of catastrophic and non-catastrophic transitions

We used the Early Warning Signal toolbox (http://www.early-warning-signals.org/) to calculate two EWS, variance and lag-1 autocorrelation, from our simulated time series. For each time series, we assessed whether the EWS rose in advance of the transition using Kendall’s-*τ*, which measures the rank correlation between EWS values and time (as in Dakos et al 2012a). We rejected the null hypothesis of no EWS rise if the computed Kendall’s-*τ* statistic for a particular simulated time series had a *p*-value ≤ 0.05. EWS should be interpreted as signaling an impending transition if *both* variance and autocorrelation rise (Ditlevsen and Johnsen 2010). For model [**6**], there is no transition and thus no natural point in the time series for us to check for a rise in EWS. For this model, we checked for EWS rise before an arbitrarily chosen time point midway through the simulated time series.

The variance and autocorrelation are calculated within a rolling window, and we repeated the EWS calculation for 3 different window sizes with lengths equal to 30%, 50%, 70% the size of the available time series. We also considered 3 different time series lengths: the longest possible interval before a transition, and intervals equal to one half and one quarter that length. The Early Warning Signals toolbox uses Gaussian kernel smoothing to detrend the time series, and we adjusted the degree of smoothing by considering 3 different bandwidths (5, 30, and 60). For all analyses, we used time series that consisted of 1 observation per time unit (*t* = 0, 1, 2, 3,…).

## Results

### EWS for models with colored noise (τ > 0)

When transitions were due to changes in the control parameters, our models varied greatly in their propensity to show EWS before a shift. Variance and autocorrelation both rose consistently in advance of the saddle node bifurcation (model [**1**]), the bifurcation that lacked CSD (model [**5**]), and often, but not always, the sharp transition without a bifurcation (model [**7**]) (figure 2). Autocorrelation, but not variance, consistently rose in model [**2**], with the transcritical bifurcation, and the opposite was true of model [**3**] (super-critical Hopf). In all other cases, EWS rose in half or less of the simulated *σ*, *τ* combinations. As expected, this includes our null case (model [**6**]) and all cases of stochastic switching (figure 2).

**Figure 2.**
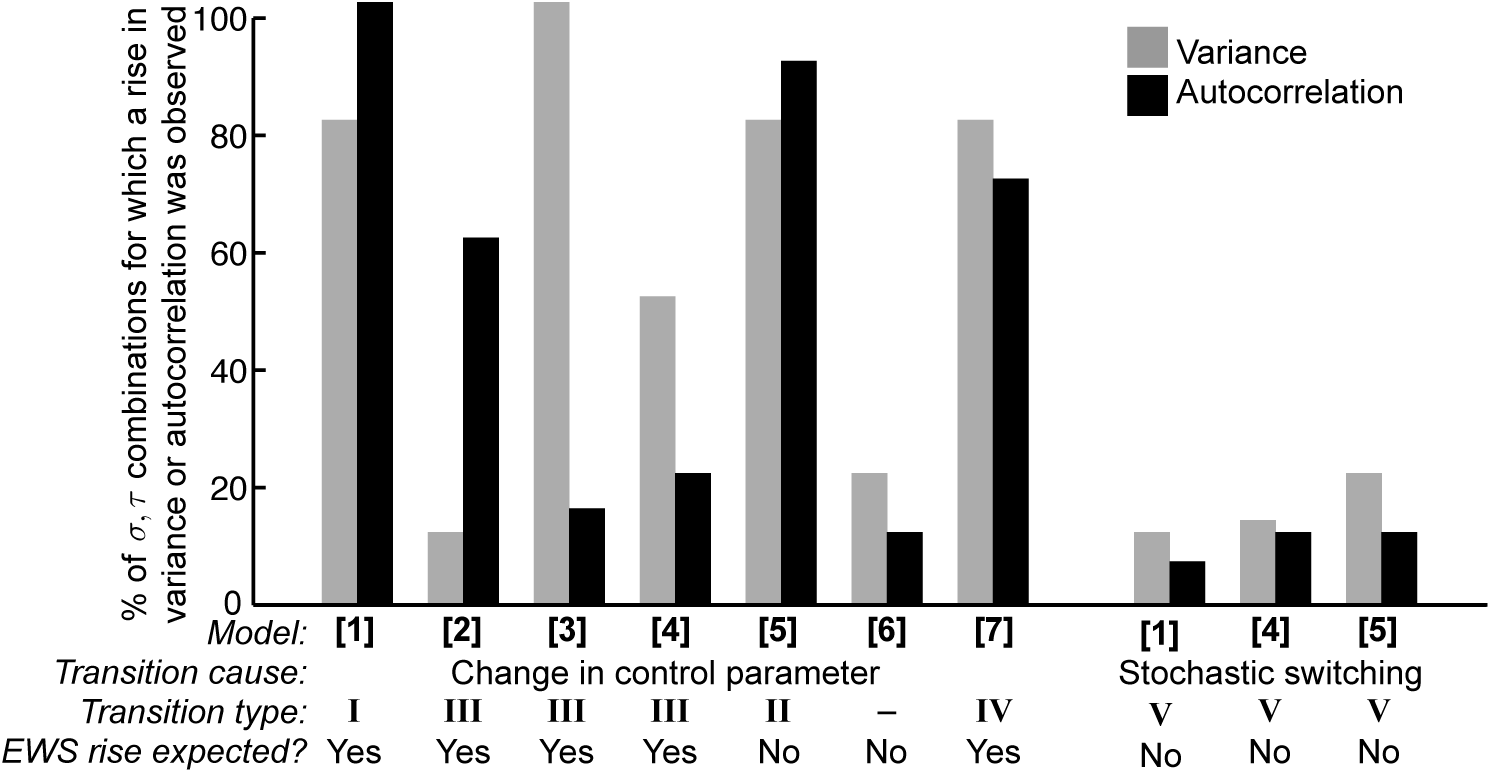
Histograms showing the percentage of colored noise simulations that showed a rise in variance or autocorrelation before a transition. Each model was simulated using the parameter values shown in table 1 across many combinations of *τ* and *σ*, as explained in the text. EWS were calculated using the longest possible pre-transition time series, a rolling window size of 50% the time series length, and a smoothing bandwidth of 40.

The method of calculating EWS had some impact on our results (figures 3-4, table 3). In particular, we found that EWS generally rose more reliably in advance of a transition when we used a longer time series leading up to that transition (figure 3c-d). Rolling window size (figure 4a) and smoothing bandwidth (figure 4b) had less effect on the EWS (table 3) in comparison with the time series length, across the ranges we considered.

**Figure 3.**
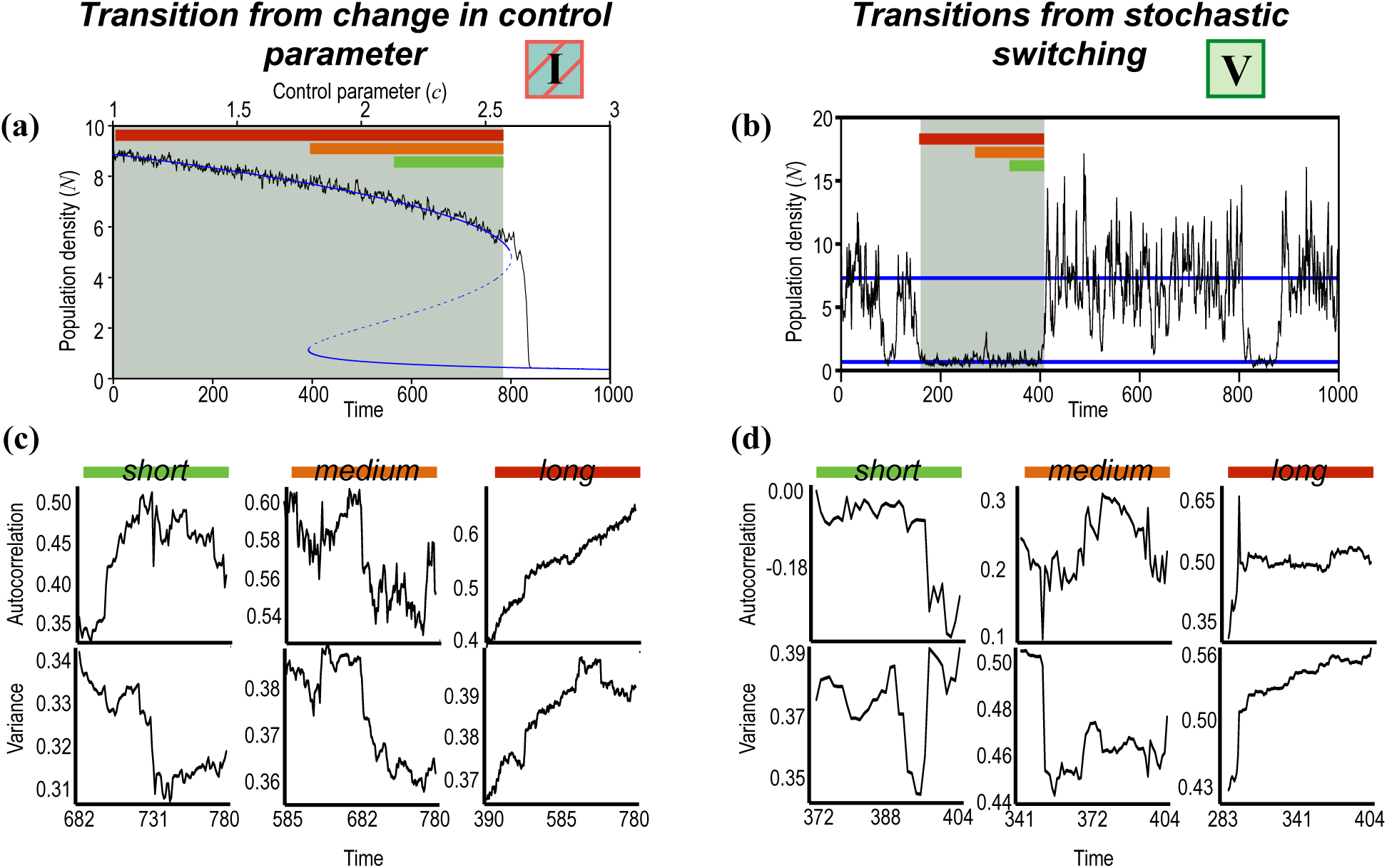
EWS for model [**1**]. The left column (a,c) considers transitions that are driven by change in the control parameter, and the right column (b,d) considers stochastic switching. (a-b): Simulated time series (black lines) plotted with stable (solid blue lines) and unstable (dashed blue lines) equilibria. The gray shaded area marks the longest time series that is available for analysis preceding a shift. (c-d): EWS calculated from 3 different time series lengths, equal to the last 25%, 50%, or 100% of the gray shaded time series (spans marked, respectively, with green, orange, and red bars in (a-b)). All model parameters are as shown in tables 1-2; (c-d) use a window size of 50% the time series length and bandwidth 5.

**Figure 4.**
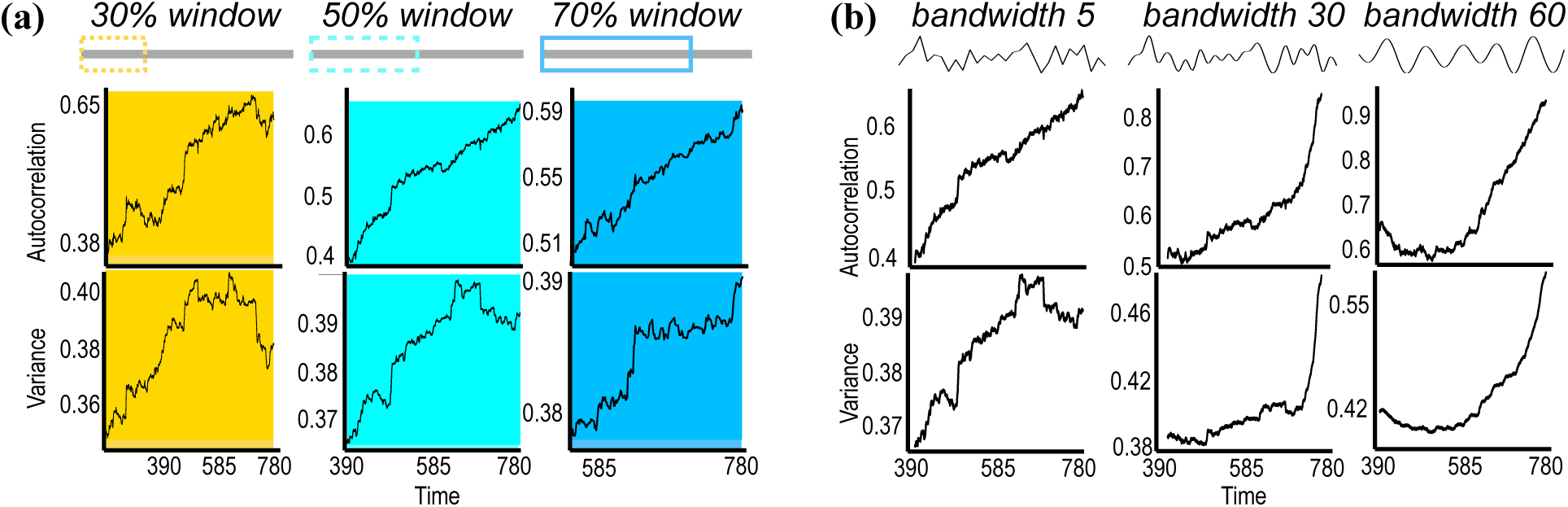
Sensitivity of EWS to window size and smoothing bandwidth for model [**1**] with a change in the control parameter (time series shown in figure 3a). (a) EWS calculated with 3 different rolling window sizes, of lengths equal to 30%, 50%, and 70% the length of the time series being analyzed. (b) EWS calculated using 3 different bandwidths: 5, 30, and 60. All model parameters are as shown in tables 1-2. The longest time series available (entire shaded region in figure 3a) was used; (a) uses bandwidth 5 and (b) uses a window size of 50% the time series length.

**Table 3.**
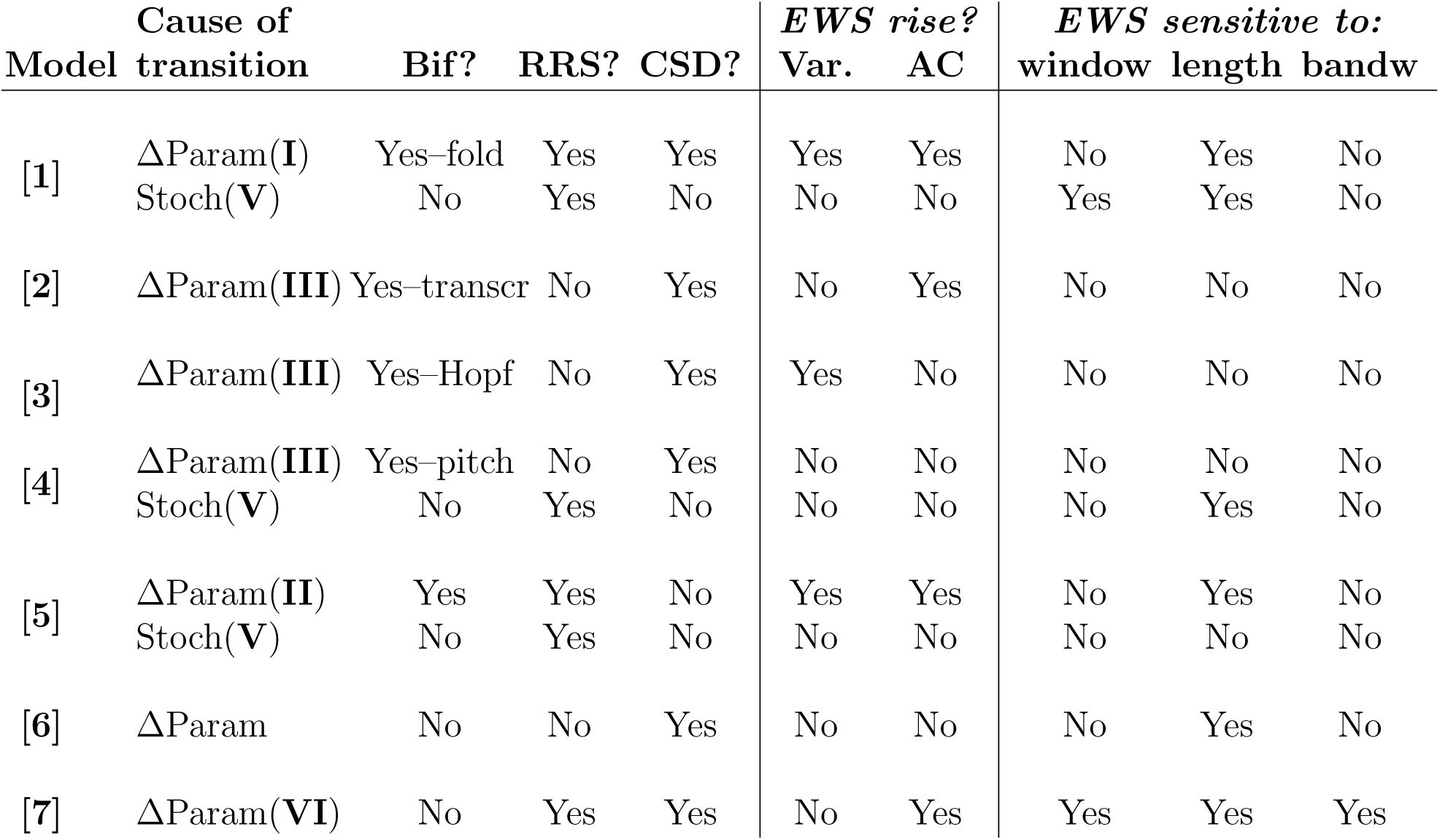
Summary of results under fixed values of *σ* and *τ* (right columns of table 2). Model equations can be found in Table 1. For each model, there are up to 2 possible causes for transitions: a change in parameter values (ΔParam) or stochastic switching (Stoch). For each transition, we list the classification according to figure 1a and specify which phenomena are present (Bif = bifurcation; RRS = rapid regime shifts; CSD = critical slowing down). We state whether the early warning signals (EWS: variance (Var.) and lag-1 autocorrelation (AC)) consistently rises in advance of the transition, based on analysis of the longest possible pre-shift time series, a rolling window of 50% the time series length, and a bandwidth of 5. Lastly, we report whether these are sensitive to window size (window), time series length (length), or bandwidth (bandw). Other abbreviations: transcr = transcritical, pitch = pitchfork.

Examples (based on parameter values shown in tables 1–3) of the simulated time series and the behavior of the EWS are shown in figures 3-5 and summarized in table 3. These examples again show clear rises in both EWS for models [**1**] and [**5**], and mixed results for the other models (including, for this set of *σ* and *τ* values, model [**7**]).

**Figure 5.**
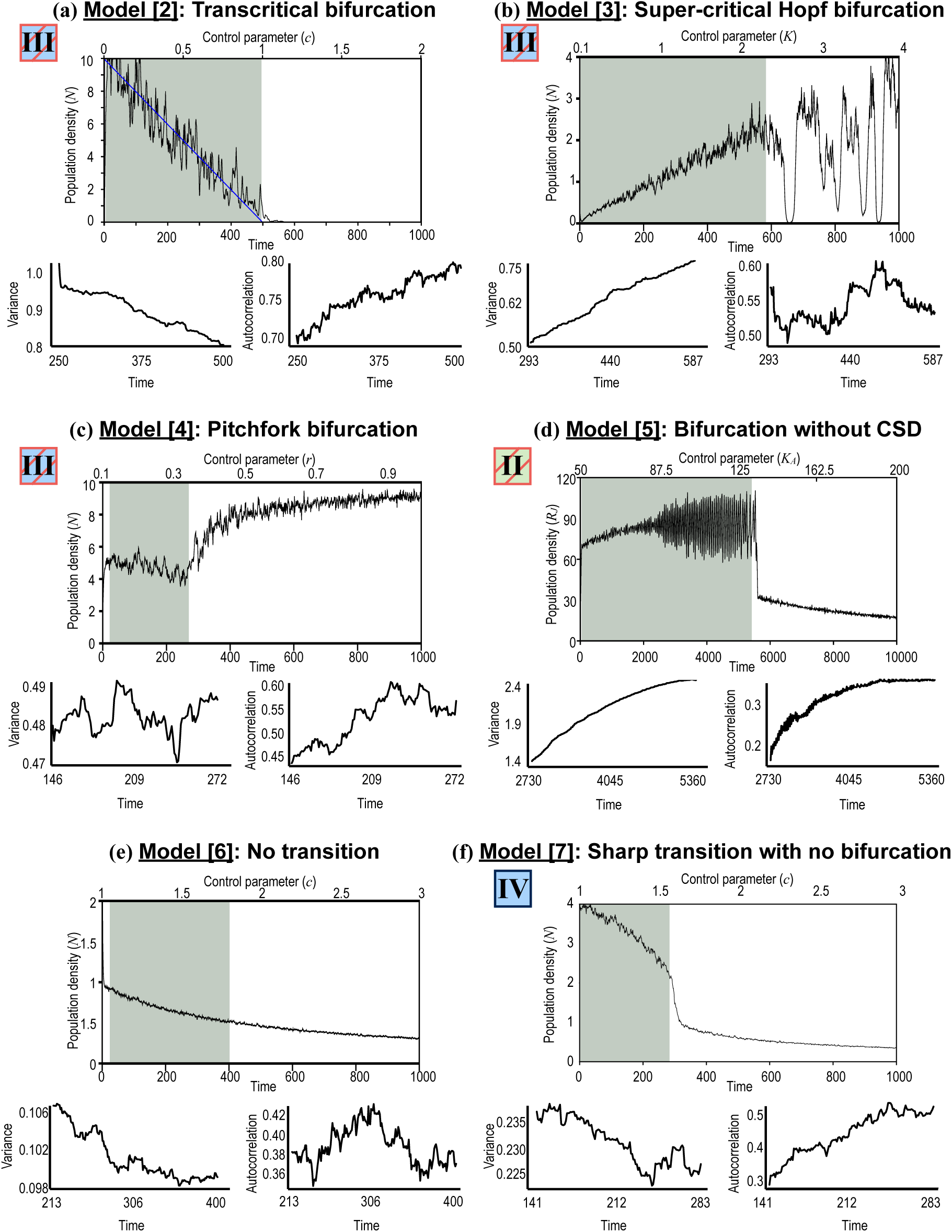
EWS for models [**2**]-[**7**] in cases where transitions are driven by changes in the control parameters. The gray shaded area shows the time series analyzed, and all results shown here used a rolling window of 50% the time series length and a bandwidth of 5. All model parameters are as shown in tables 1-2.

### EWS for models with white noise (*τ* → 0)

When we simulated each model with multiplicative white noise (equations (1)-(2) with *τ* → 0), both variance and autocorrelation usually rose in advance of a transition caused by a parameter change (figure 6). This was true regardless of whether CSD was present (in which case we expect both signals to rise; models [**1**]-[**4**], [**7**]) or not (model [**5**]). Our null model [**6**] showed a rise in both signals, and especially variance, in a surprisingly large number of cases, give that this model has no transition. EWS again typically did not rise before stochastic switching (figure 6). Figure 7 provides a comparison of our results (for multiplicative red and multiplicative white noise) and those of Kéfi et al (2012) (for additive white noise).

**Figure 6.**
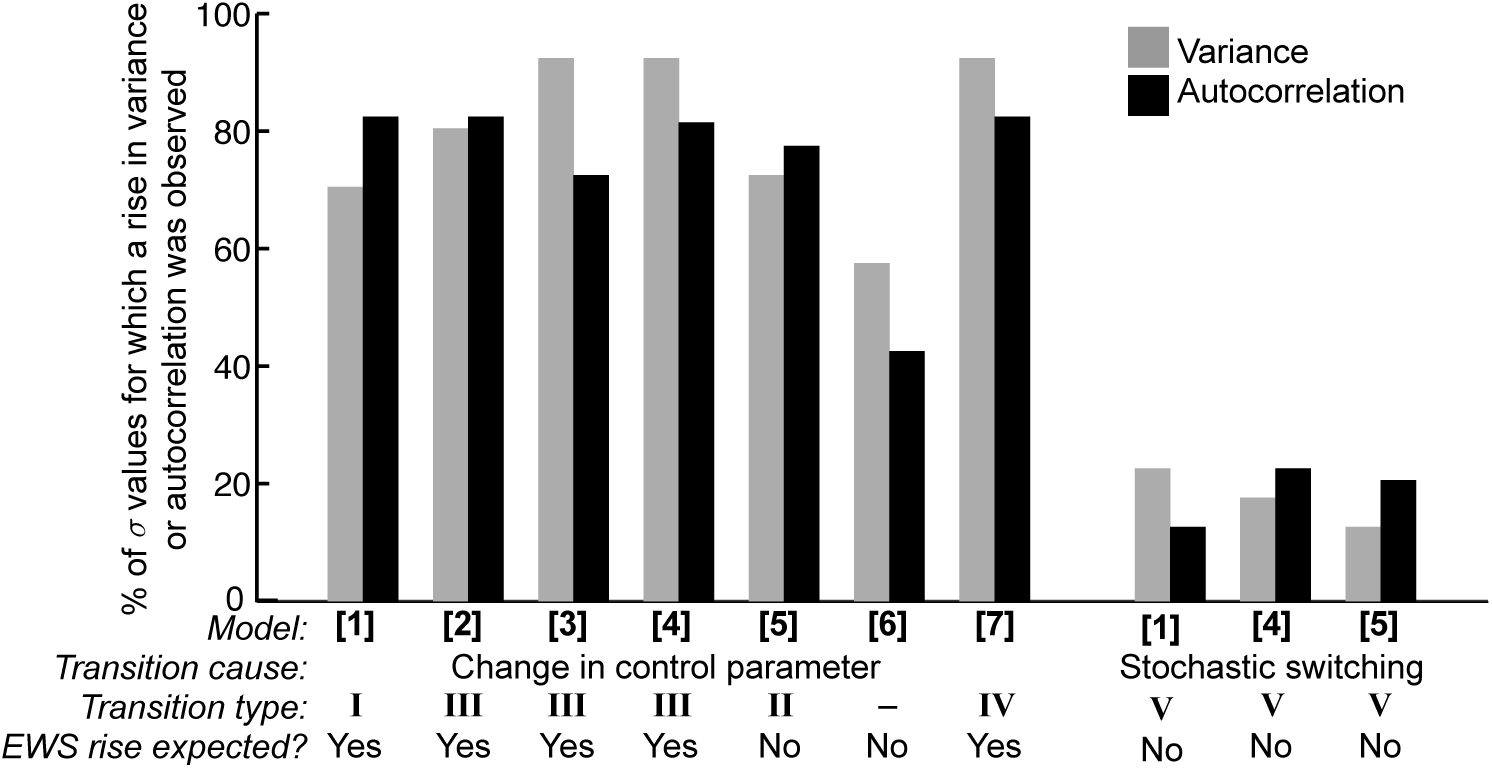
Histograms showing the percentage of simulations that showed a rise in variance or autocorrelation before a transition when models were simulated under multiplicative white noise. Parameter values as in figure 2.

**Figure 7.**
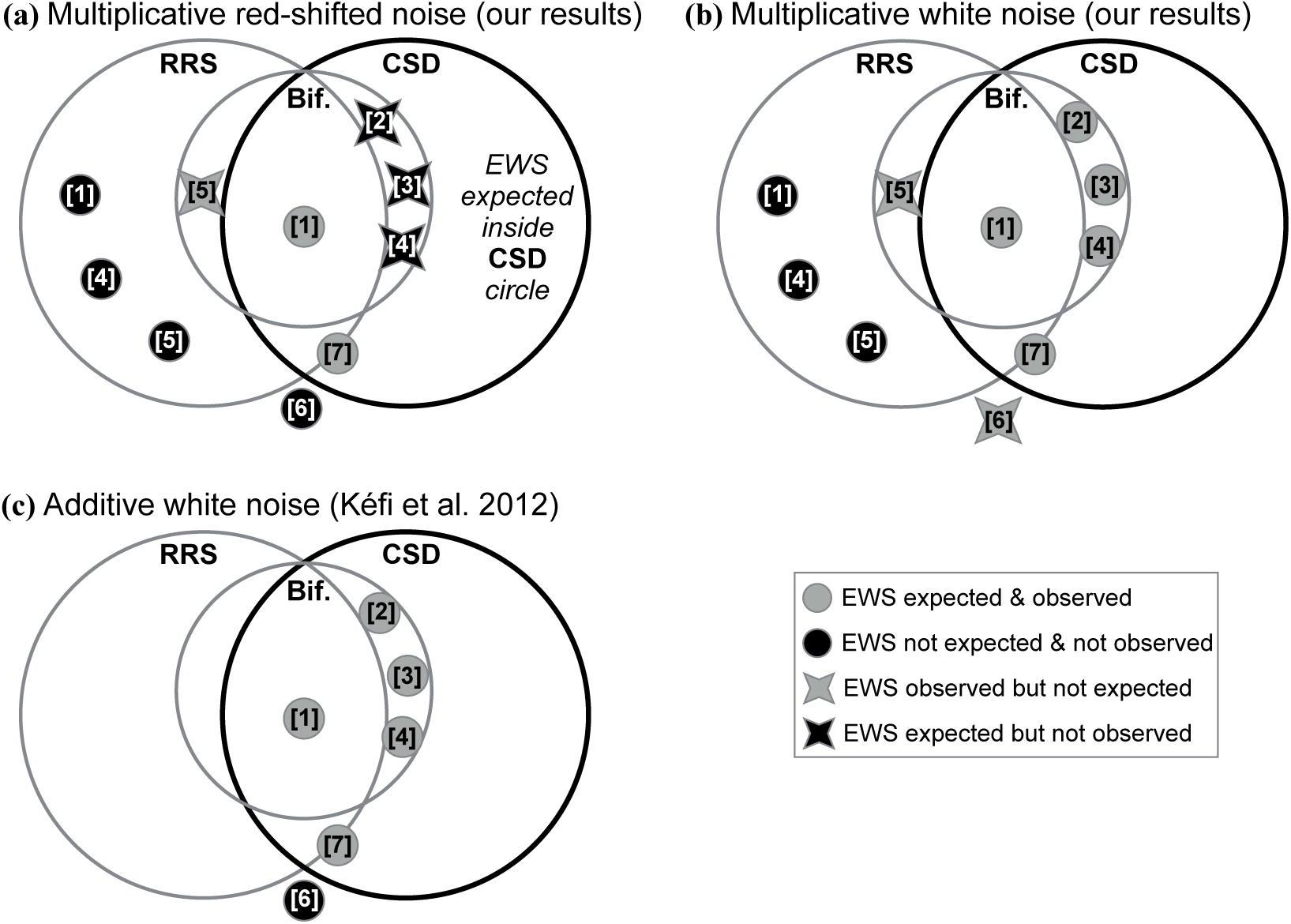
Graphical summary of all results, using the same Venn diagram as in figure 1a. (a) Our results using multiplicative colored noise; (b) our results using multiplicative white noise; (c) results reported for additive white noise in Kéfi et al (2012). For the purposes of summarizing our results, we say “EWS observed” if both variance and autocorrelation rose in ≥ 40% of *σ*, *τ* combinations (panel a) or *σ* values (panel b) that we examined. Numbers in square brackets refer to model numbers. Models numbers inside dots behaved as expected: both EWS consistently rose in advance of a transition with CSD (“EWS expected”) or failed to rise in advance of a transition without CSD (“EWS not expected”). Models inside X-shaped symbols did not behave as expected. Gray shapes mark all cases where a rise in both variance and autocorrelation was observed, and black shapes mark cases where one or both signals did not consistently rise. If the EWS only behaved as expected, the CSD circle would contain only gray dots, and all shapes outside this circle would be black dots. Note that Kéfi et al (2012) did not consider our model [**5**] nor any cases of stochastic switching, so those points are not depicted in (c). Note also that while Kéfi et al (2012) did report EWS in model [**6**], they observed the EWS rise outside the range of the control parameter that we used here; therefore, we mark [**6**] as “EWS not observed” in (c) for a proper comparison against our results.

## Discussion

Our analyses showed both surprising positive and surprising negative results. With multiplicative red-shifted noise, the early warning signals appeared in advance of a transition without critical slowing down (model [**5**]) and failed to appear in advance of several transitions with CSD (models [**2**], [**3**] and [**4**]) (figure 7a). When we instead modeled multiplicative white noise, the EWS rose as expected before all transitions with CSD, but still frequently also rose before a non-CSD transition (again model [**5**]) and before an arbitrarily-chose time point in the model that lacked a transition ([**6**]) (figure 7b). Our results for using multiplicative white noise are in close agreement with Kéfi et al’s (2012) results with additive white noise (figure 7c). Agreement between EWS in the presence of additive and multiplicative noise was also reported by Kuehn (2013). It therefore appears that the performance of EWS is more sensitive to noise color (temporal autorcorrelation) than to the exact way stochasticity enters into the model.

Encouragingly, we found that EWS are robust indicators of an impending saddle-node (fold) bifurcation regardless of the type of noise we used. In some sense, this is unsurprising: saddle-node bifurcations fit within region I (figure 1a), which Boettiger et al (2013) referred to as the “charted territory” because the vast majority of EWS research in ecology has been conducted on models in this region. Nevertheless, even in region I, there is a general paucity of ecological studies with non-additive noise (Hastings and Wysham 2010), and past reports on the effect of noise color on EWS have been mixed (Rudnick and Davis 2003, Dakos et al 2012b, Perretti and Munch 2012, Boerlijst et al 2013). EWS can fail to predict an approaching fold in some instances of anisotropic perturbations (i.e. noise that affects some components of the system more strongly than others; Boerlijst et al 2013). We are not, therefore, suggesting that EWS will universally warn of fold bifurcations. Still, we find it encouraging and interesting that they worked so well under multiplicative red and white noise. While red-shifted noise obviously affects population autocorrelation, it does not appear to interfere with the *trend* in autocorrelation that signals an approaching transition due to a fold bifurcation.

In contrast, these EWS appear unreliable in the context of other kinds of transitions. We expected the EWS to rise in advance of any transition with CSD but for models [**2**]-[**4**], we only saw this rise for white noise. Red-shifted noise did appear to interfere with the use of autocorrelation as an EWS for models [**3**] and [**4**] (see “change in control parameter” bars, figure 2 (red noise) versus figure 6 (white noise)). However, for model [**2**], and to a lesser extent [**4**], it was variance that failed to rise under red noise. When CSD was absent, and so EWS were not expected to rise, they still consistently rose for some transitions. There are two important lessons in these observations. First, EWS can rise before some transitions (or arbitrary moments, for our null model without a transition) that lack CSD. This highlights the plain fact that properties like variance and autocorrelation have many causes unrelated to critical slowing down, and that their patterns are not always driven by critical points. Second, EWS do not appear to be robust indicators of CSD in the uncharted territories of III and IV (where, of course, even if CSD is detected, it will not (III) or may not (IV) be associated with a rapid regime shift at the bifurcation point). We therefore strongly second Boettiger et al’s (2013) remark that “establishing the saddle-node mechanism is a necessary condition of using [**the EWS of**] CSD as a warning signal.” Developing strategies for identifying impending dynamical changes other than the saddle-node bifurcation is an exciting open challenge.

We placed stochastic switching into category V (figure 1a), as an example of a rapid regime shift that is not accompanied by a bifurcation nor CSD. Boettiger et al (2013) also used this classification, but there has been some debate about whether slowing down should be expected in some cases of stochastic switching (Boettiger and Hastings 2013, Drake 2013). Drake (2013) argued that when there is a shift between stable states separated by an unstable equilibrium, CSD should be observed as the system traverses the unstable point (at which the eigenvalue equals zero). However, we found no EWS before a stochastic switch, regardless of whether that switch crossed an intervening unstable point (models [**1**] and [**4**]) or not (model [**5**]). We instead found support for the idea that unstable states are traversed sufficiently quickly that no CSD appears, in agreement with theory (Boettiger and Hastings 2013, Freidlin and Wentzell 2012).

We found, like Dakos et al (2012a), that variance and autocorrelation are typically robust to the rolling window size and smoothing bandwidth used to compute them (table 3). Both indicators were much more sensitive to the length of the pre-transition time series analyzed. In most cases, using a longer time series improved EWS performance (i.e. made them more likely to rise or not as expected). However, for model [**5**]’s transition due to changing the control parameter, the unexpected rise in variance and autocorrelation was *only* observed in the longest time series. Thus, for model [**5**] the EWS would have behaved as expected if we had used less pre-transition data. The null model [**6**] behaved as expected for the longest and shortest time series considered, but not when we used an intermediate length. Together, these results suggest that having more data does not invariably reduce the unexpected behaviors of EWS. Even – and in some cases, especially – with long datasets, calculating EWS without first knowing that a saddle-node bifurcation exists in the system can lead to faulty conclusions.

We conclude that EWS performance is robust to differences in noise type in the case of the saddle-node bifurcation, for the models and (isotropic) noise types we considered. For other transitions, though, EWS did not necessarily accompany CSD, and their behavior was highly sensitive to noise type. Continued research on the development and refinement of early warning signals has rightly garnered much attention (Scheffer et al 2009, Dakos et al 2012a,b); the prospect of being able to predict a catastrophic shift before it occurs is truly exciting. At the same time, any early warning signal will have bounds on its applicability (Kéfi et al 2012, Boettiger and Hastings 2012b, Dakos et al 2015, Gsell et al 2016). Research the helps us define and understand these boundaries improves our ability to properly recognize warning signals and points the way toward open areas in need of new EWS development.

## Acknowledgements

PSD acknowledges financial support from ISIRD, IIT Ropar Grant No.: IITRPR/Acad./52. This work was partially supported by a Complex Systems Scholar grant to KCA from the James S. McDonnell Foundation. We thank the members of the Abbott and Snyder labs at Case Western Reserve University for extremely helpful suggestions on presentation.

